# Neonatal hyperoxia inhibits proliferation of atrial cardiomyocytes by suppressing fatty acid synthesis

**DOI:** 10.1101/2020.06.01.127621

**Authors:** Ethan David Cohen, Min Yee, George A. Porter, Andrew N. McDavid, Paul S. Brookes, Gloria S. Pryhuber, Michael A. O’Reilly

**Author notes:** **Address Correspondence to:** Ethan David Cohen, Ph.D., Department of Pediatrics, Box 850, The University of Rochester, School of Medicine and Dentistry, 601 Elmwood Avenue, Rochester, NY 14642, Michael A. O’Reilly, Ph.D., Department of Pediatrics, Box 850, The University of Rochester, School of Medicine and Dentistry, 601 Elmwood Avenue, Rochester, NY 14642.

## Abstract

Preterm birth increases the risk for pulmonary hypertension and heart failure in adulthood. Oxygen therapy can damage the immature cardiopulmonary system and may be partially responsible for the cardiovascular disease in adults born preterm. We previously showed that exposing newborn mice to hyperoxia causes pulmonary hypertension by 1 year of age that is preceded by a poorly understood loss of pulmonary vein cardiomyocyte proliferation. We now show that hyperoxia also inhibits the proliferation of left atrial cardiomyocytes and causes diastolic heart failure by thinning the walls of the left atrium and disrupting its ability to pump effectively. Transcriptomic profiling showed that neonatal hyperoxia permanently suppressed fatty acid synthase (*Fasn*), stearoyl-CoA desaturase 1 (*Scd1*) and other fatty acid synthesis genes in the atria of mice, the HL-1 line of mouse atrial cardiomyocytes and left atrial tissue explanted from human infants. Suppressing *Fasn* or *Scd1* reduced HL-1 cell proliferation while overexpressing these genes maintained their expansion in hyperoxic conditions, suggesting hyperoxia directly inhibits atrial cardiomyocyte proliferation by repressing *Fasn* and *Scd1*. Pharmacologic interventions that restore *Fasn, Scd1* and other fatty acid synthesis genes in atrial cardiomyocytes may thus provide a way of ameliorating the adverse effects of supplemental oxygen on preterm infants.

## INTRODUCTION

Approximately 10% of births occur before 37 weeks of gestation and are thus considered preterm. Children who are born preterm face an increased risk of developing airway hyperreactivity, chronic wheezing, emphysema and other respiratory diseases as children and young adults than those born at term (1–5). Now that more and more severely preterm infants are reaching adulthood, it is becoming clear that these individuals are also predisposed to pulmonary vascular disease and heart failure (6, 7). Most strikingly, people who were born extremely preterm are 4.6 times more likely to develop pulmonary hypertension and 17 times as likely to suffer heart failure as young adults than people who were born at term (8). While prior studies have reported a loss of pulmonary capillaries, altered ventricular shape and decreased aortic size in young adults born preterm (9–13), the root causes of pulmonary hypertension, heart failure and other cardiovascular diseases in the survivors of preterm birth remain poorly understood. Recent echocardiographic, magnetic resonance, and CT imaging studies indicate that preterm infants have larger left atria and lower diastolic function than term infants (14–17). These changes may be an early and direct response to oxygen exposure because diastolic heart failure has been seen in preterm infants on supplemental oxygen therapies (18).

We and other investigators have been using rodents to understand how early-life exposure to high levels of oxygen (hyperoxia) causes cardiovascular disease. In our model, newborn mice are exposed to room air or hyperoxia (100% oxygen) from birth to postnatal day (PND) 4 and then recovered in room air. Mice exposed to neonatal hyperoxia develop pulmonary diseases like those of former preterm infants, including alveolar simplification non-atopic airway hyperreactivity (19) and reduced lung function (20). They also have persistent inflammation and develop fibrotic lung disease after influenza A infection (21,22). Additionally, hyperoxia-exposed mice develop pathological symptoms of pulmonary hypertension that include pulmonary capillary rarefaction, right ventricular hypertrophy, and a 50% mortality by one year of age (23). Other investigators have similarly observed cardiovascular disease, including right ventricular hypertrophy, hypertension, capillary rarefication and other types of vascular dysfunction, in animals exposed to neonatal hyperoxia (24–28). Together, these findings demonstrate that neonatal hyperoxia can cause pulmonary hypertension and other cardiovascular diseases in adult rodents.

We recently reported that hyperoxia inhibits the proliferation of the cardiomyocytes wrapping the pulmonary vein and prevents them from expanding to cover its distal branches within the growing lungs of neonatal mice (29). Since these cells form a contractile sleeve around the pulmonary vein and extending into the left atria, the early loss of these cells may increase the force needed to drive pulmonary circulation and thus contribute to the development of pulmonary venous congestion and heart failure in aged hyperoxia-exposed mice (30). Herein, we report that neonatal hyperoxia inhibits the proliferation of left atrial cardiomyocytes by suppressing genes required for the *de novo* synthesis of fatty acids. Failure to expand these cells results in hypoplastic left atria that dilates as mice age and eventually lose the ability to effectively fill the left ventricle, a finding that may help explain the pathogenesis of diastolic heart failure in the survivors of preterm birth.

## RESULTS

### Neonatal hyperoxia inhibits cardiomyocyte proliferation in the left atrium and causes diastolic heart failure in adult mice

We previously showed that hyperoxia causes a loss of the cardiomyocytes wrapping the pulmonary vein by inhibiting their proliferation (29). Since the pulmonary vein cardiomyocytes are contiguous with the left atria, we investigated whether hyperoxia also inhibited proliferation of left atrial cardiomyocytes. H&E staining revealed that the left atria of PND4 mice exposed to hyperoxia were larger than controls (dotted lines, **Figure 1A, B**). Staining for the mitosis marker phosphorylated histone H3 (pHH3) and the cardiomyocyte marker cardiac troponin T (TNNT2) showed that there were fewer proliferating cardiomyocytes in the left atria of hyperoxia-exposed mice than in controls (arrows, **Figure 1C, D**). The density of nuclei in TNNT2+ areas of the left atria were also lower in hyperoxia-exposed mice than in controls (**Figure 1E**), indicating the left atrial cardiomyocytes underwent hypertrophy in order to compensate for fewer cells.

**Figure 1.**
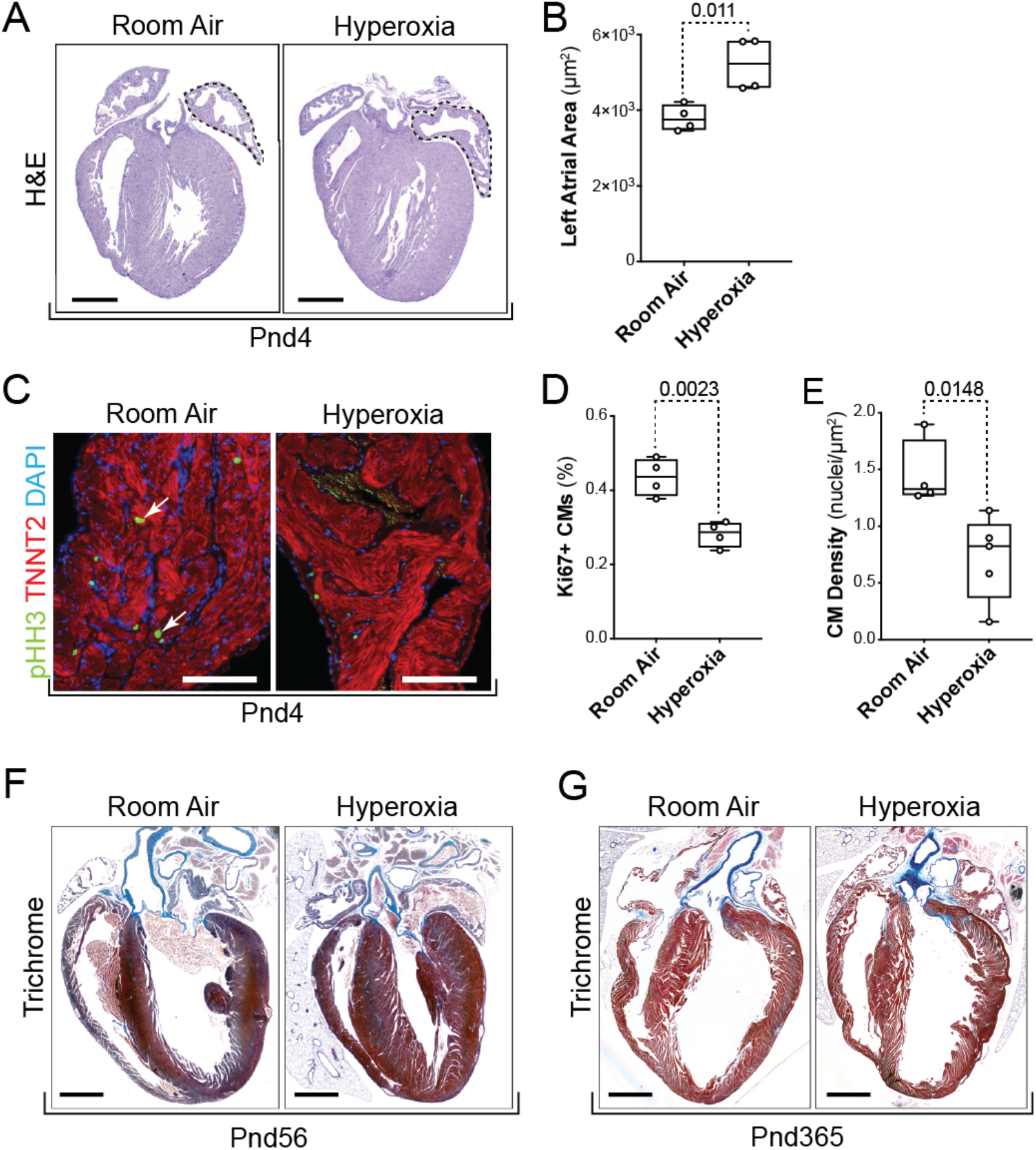
Neonatal hyperoxia inhibits left atrial cardiomyocyte (CM) proliferation. (A) Hematoxylin and eosin (H&E) stained sections of hearts harvested from postnatal day (PND) 4 mice that were exposed to room air (left) or hyperoxia (right). Dotted lines outline the left atria. Scale bar = 200μm (B) Mean area of the left atria in sections of room air and hyperoxia exposed mice on PND4. (C) Sections of left atrial appendages from PND4 control (left) and hyperoxia-exposed (right) mice stained with phosphorylated Histone H3 (pHH3, green), cardiac troponin T (TNNT2, red) and 4’,6-diamidino-2-phenylindole (DAPI, blue). Arrows point to pHH3+ cardiomyocytes. Scale bars = 100μm (D) Percentage of pHH3+ cardiomyocytes in PND4 mice exposed to room air or hyperoxia. (E) Numbers of DAPI stained nuclei per mm^2^ of TNNT2 labeled myocardium. (F, G) Sections of hearts from mice exposed to room air (left) or hyperoxia (right) from PND0-4 and then returned to room air until PND56 (F) or PND365 (G). Arrows point to the dilated left atria of hyperoxia-exposed mice. Scale bars = 400μm (B, D, E) Box plots show the median, second and third quartiles of measurements from 4 room air and 4 hyperoxia-exposed mice, whiskers indicate the range and circles represent individual data points. The p-values between datasets are from unpaired t-tests.

To determine how the loss of left atrial cardiomyocyte proliferation affects adult cardiac morphology, hearts were collected from PND56 (**Figure 1F**) and PND365 (**Figure 1G**) mice who had been exposed to room air or hyperoxia between PND0-4. The left atria of hyperoxia-exposed mice remained enlarged relative to control, but had thinner walls suggesting that the initial hypertrophic response to hyperoxia progressed to dilated cardiomyopathy. In contrast, the ventricles and right atria of PND56 mice exposed to neonatal hyperoxia were mildly thickened relative to controls. Interestingly, the ventricular walls of hyperoxia-exposed mice were no longer thickened on PND365. However, the right ventricles of hyperoxia exposed mice were enlarged relative to controls, consistent with our prior finding that neonatal hyperoxia increased right ventricular mass by this age (23).

To determine how neonatal hyperoxia affects adult heart function, echocardiography was performed on PND56 and PND365 mice who had been exposed to room air or hyperoxia between PND0-4 (**Supplemental Table 1**). Despite the increased left ventricular mass observed in hyperoxia-exposed mice on PND56, cardiac function was not different than controls at this age. In contrast, significant changes in cardiac function were seen in PND365 mice exposed to neonatal hyperoxia. The left ventricles of hyperoxia-exposed mice harvested on PND356 were smaller than controls during diastole but not during systole (**Figure 2A, B, C, D**). The stroke volume of the left ventricle was thus reduced in mice exposed to neonatal hyperoxia relative to controls (**Figure 2E**). While the heart rates of mice were unaffected, neonatal hyperoxia reduced the stroke volumes and cardiac outputs of mice (**Figure 2F, G**). The left ventricular ejection fractions of mice exposed to neonatal hyperoxia tended to be lower than those of controls but this difference was relatively small and failed to reach statistical significance (**Figure 2H**). Ratios of the early velocity of left ventricular filling to the early velocity of flow past the mitral valve annulus (E/e’), which is used to assess the early diastolic filling of the left ventricle as it relaxes, was unchanged in mice exposed to neonatal hyperoxia (data not shown). The reduced filling of the left ventricle in mice exposed to neonatal hyperoxia is thus likely to be caused by either defective venous return from the lungs or reduced left atrial contractility and not from stiffening of the left ventricular myocardium due to pathological fibrosis.

**Figure 2.**
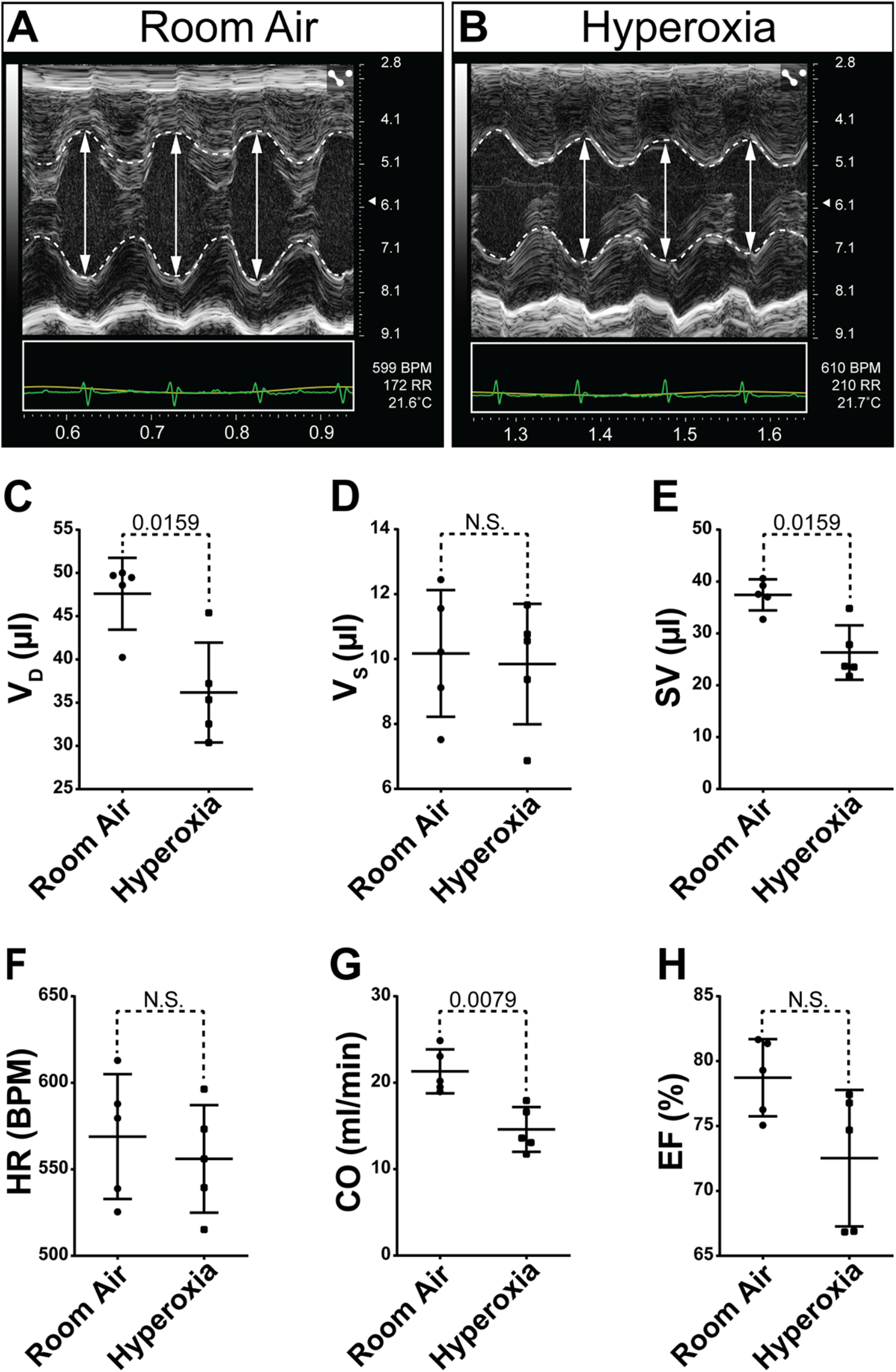
Neonatal hyperoxia causes mice to develop dilated heart failure later in life. (A, B) Short axis M-mode echocardiograms of PND365 mice exposed to room air (A) or hyperoxia (B) from birth to PND4. Dotted lines trace the walls of the left ventricle (LV). Double headed arrows show the diameters of the LV in diastole. (C-H) Graphs show the volume in diastole (V_D_, C), volume in systole (V_S_, D), stroke volume (SV, E), heart rate (HR, F), cardiac output (CO, G) and ejection fraction (EF, H) the LVs of neonatal hyperoxia-exposed and control mice on PND365. Bars mark the mean and 95% CI for measurements from 5 neonatal hyperoxia-exposed and 5 control mice. Dots show single data-points, p-values from Mann-Whitney tests are shown between datasets.

### Neonatal hyperoxia inhibits the *de novo* fatty acid synthesis pathway in the atria of mice

An Affymetrix array was used to identify hyperoxia-induced changes in gene expression responsible for the inhibition of left atrial cardiomyocyte proliferation. RNA was isolated from the atria of 4 individual hyperoxia-exposed and 3 room air control mice on PND4 and hybridized to the array. Out of 39,000 transcripts examined, only 158 differed by ≥ 1.5-fold with a p-value < 0.05 and false discovery rate < 0.3 (**Figure 3A**). Neonatal hyperoxia increased expression of 53 genes (**Supplemental Table 2**) and inhibited expression of 105 genes (**Supplemental Table 3**). Gene ontology (GO) analysis identified several overlapping sets of genes involved in small molecule biosynthetic processes, monocarboxylic acid metabolism, lipid metabolic processes and the regulation of lipid metabolism (**Figure 3B**). Among the 12 genes involved in fatty acid metabolism, 4 were upregulated in the atria of hyperoxia-exposed mice relative to controls, including peroxisome proliferator-activated receptor gamma coactivator 1-alpha (*PPARGC-1α*), a transcriptional co-regulator that binds to peroxisome proliferator-activating receptor a and other transcription factors to promote mitochondrial biogenesis (31). Hyperoxia also upregulated hydroxyacyl-CoA dehydrogenase trifunctional multienzyme complex subunit alpha (*Hadha*), part of the mitochondrial complex that catalyzes the β-oxidation of long-chain fatty acids (32), and lipoprotein lipase (*Lpl*), which cleaves lipids from dietary lipoproteins (33).

**Figure 3.**
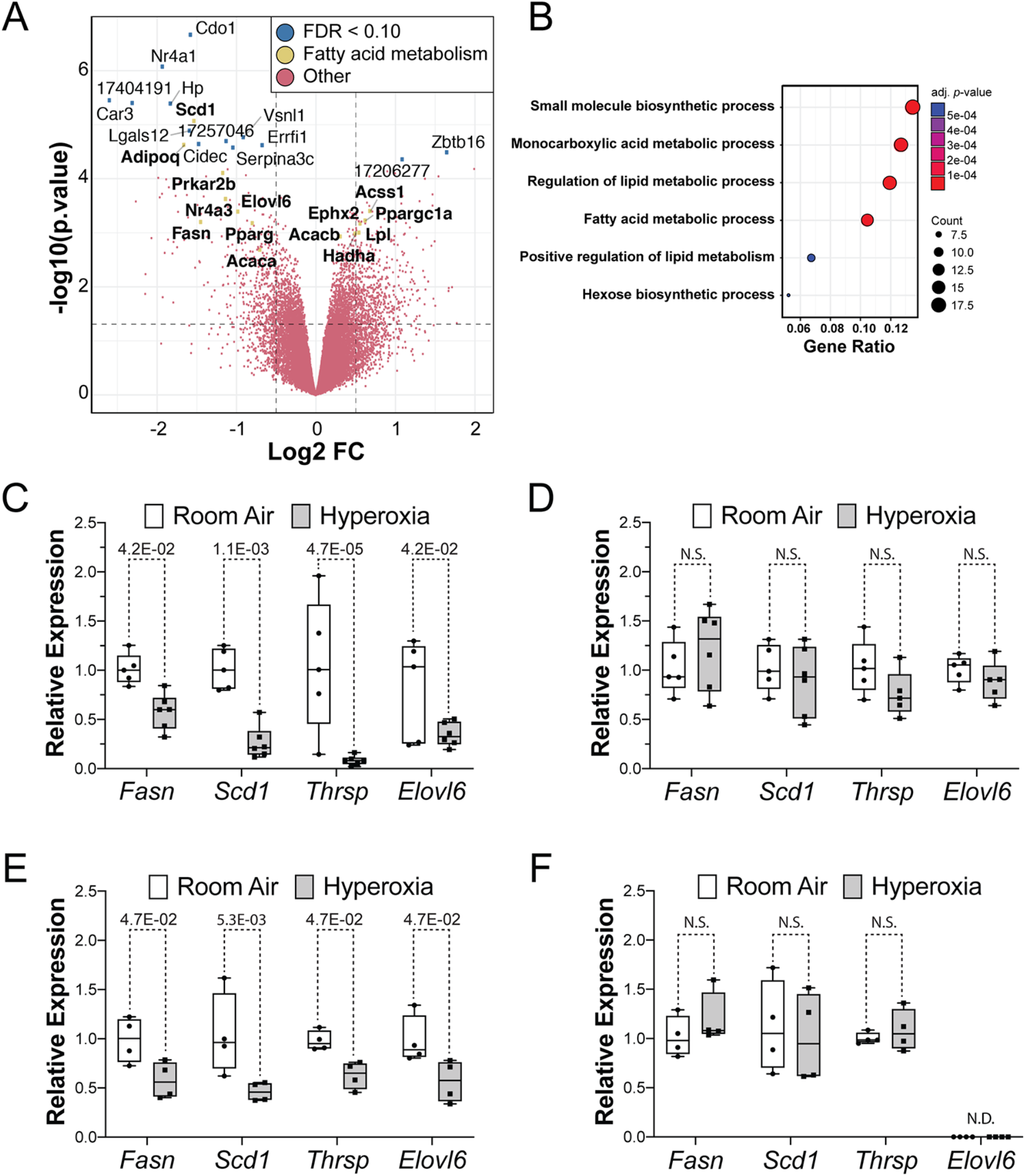
Neonatal hyperoxia suppresses genes needed for *de novo* fatty acid synthesis in atria of mice. (A) Volcano plot of Log2 fold changes vs log10 p values for differential expression of genes. Genes with FDR < 10% or annotation in the fatty acid pathway are highlighted. Genes with false discovery rate <0.1 are marked by blue dots, genes involved in fatty acid metabolism are marked by green dots and written in bold type. (B) Gene ontology (GO) analysis of differentially expressed genes. (C-F) Results of qRT-PCR for *Fasn, Scd1, Thrsp* and *Elovl6* in the atria (C, E) and ventricles (D, E) of neonatal hyperoxia-exposed and control mice on PND4 (C, D) and PND56 (E, F). Box plots show median values and inner quartiles. Error bars indicate the range of values. Filled circles and squares mark values for individual room air and hyperoxia-exposed mice, respectively. The p-values from multiple t-test with Holmes-Sidak correction are shown between conditions. (C, D) n = 5 room air and 6 hyperoxia-exposed mice. (E, F) n = 4 room air and 4 hyperoxia exposed mice.

Neonatal hyperoxia reduced the expression of 8 genes involved in fatty acid metabolism including the core enzymes in the *de novo* fatty acid synthesis pathway such as *fatty acid synthase (Fasn*), the enzyme that synthesizes the 16-carbon saturated fatty acid palmitate from acetyl-CoA and malonyl-CoA (34); *elongation of very long chain fatty acids 6 (Elovl6*), the enzyme that extends palmitate to produce the 18 carbon saturated fatty acid stearate (35); and *stearoyl-CoA desaturase 1 (Scd1*), which convert palmitate and stearate to the mono-unsaturated fatty acids palmitoleate and oleate (36), and *thyroid hormone-inducible hepatic protein (Thrsp*), which binds FASN and increases its activity (37). Neonatal hyperoxia also downregulated cell death inducing *DFFA like effector c (Cidec*), a protein required for the storage of triglycerides as lipid droplets (38), and *adiponectin (Adipoq*), an adipokine that protects cardiomyocytes from oxidative stress and apoptosis following ischemic injury (39).

The suppression of fatty acid synthesis genes intrigued us because cardiomyocytes use fatty acid ß-oxidation to produce the vast amounts of ATP needed for contractility, and are thus very sensitive to changes in lipid metabolism (40). While embryonic cardiomyocytes use glycolysis to produce the majority of their ATP, neonatal cardiomyocytes were shown to require β-oxidation to fuel their proliferation (41). Using an independent set of mice, quantitative reverse transcription PCR (qRT-PCR) confirmed neonatal hyperoxia suppressed expression of *Fasn, Scd1, Thrsp* and *Elovl6* on PND4 (**Figure 3C**). In contrast, *Fasn, Scd1, Thrsp,* and *Elovl6* were unaffected in the ventricles of these same mice at this time (**Figure 3D**). Since mice exposed to hyperoxia are returned to room air on PND4, we investigated when expression of these genes returned to control levels. Surprisingly, mRNA levels of *Thrsp, Fasn, Scd1,* and *Elovl6* remained suppressed on PND56 (**Figure 3E**). These changes were again specific for the atrium and the expression of *Thrsp, Fasn,* or *Scd1* in the ventricle was unaffected on PND56, while *Elovl6* was not detected (**Figure 3F**). Together, these data suggest that hyperoxia permanently suppresses fatty acid synthesis in the atria but not ventricles.

Immunohistochemistry was used to confirm that neonatal hyperoxia inhibited expression of FASN and SCD1 protein in left atria but not ventricular cardiomyocytes. Staining for FASN protein (**red, Figure 4 A, C, D, F**) and TNNT2 (**green, Figure 4B, E, C, F**) detected FASN in both the atrial (**arrows**) and ventricular (**arrowheads**) cardiomyocytes of PND4 mice exposed to room air (**Figure 4 A-C**). FASN was reduced in the left atrial cardiomyocytes of hyperoxia-exposed mice but unaffected in ventricular cardiomyocytes (**Figure 4D-F**). Intense staining for SCD1 was detected in the left atrial cardiomyocytes of mice exposed to room air (**arrows, Figure 4 G-I**) while much lower levels were observed in the left atria of neonatal mice exposed to hyperoxia (**arrows, Figure 4 J-L**). As observed for FASN, SCD1 levels were similar in the ventricles of hyperoxia-exposed and room air control mice (data not shown). Hyperoxia also reduced staining for THRSP in the left atria but not ventricles of the mice (data not shown).

**Figure 4.**
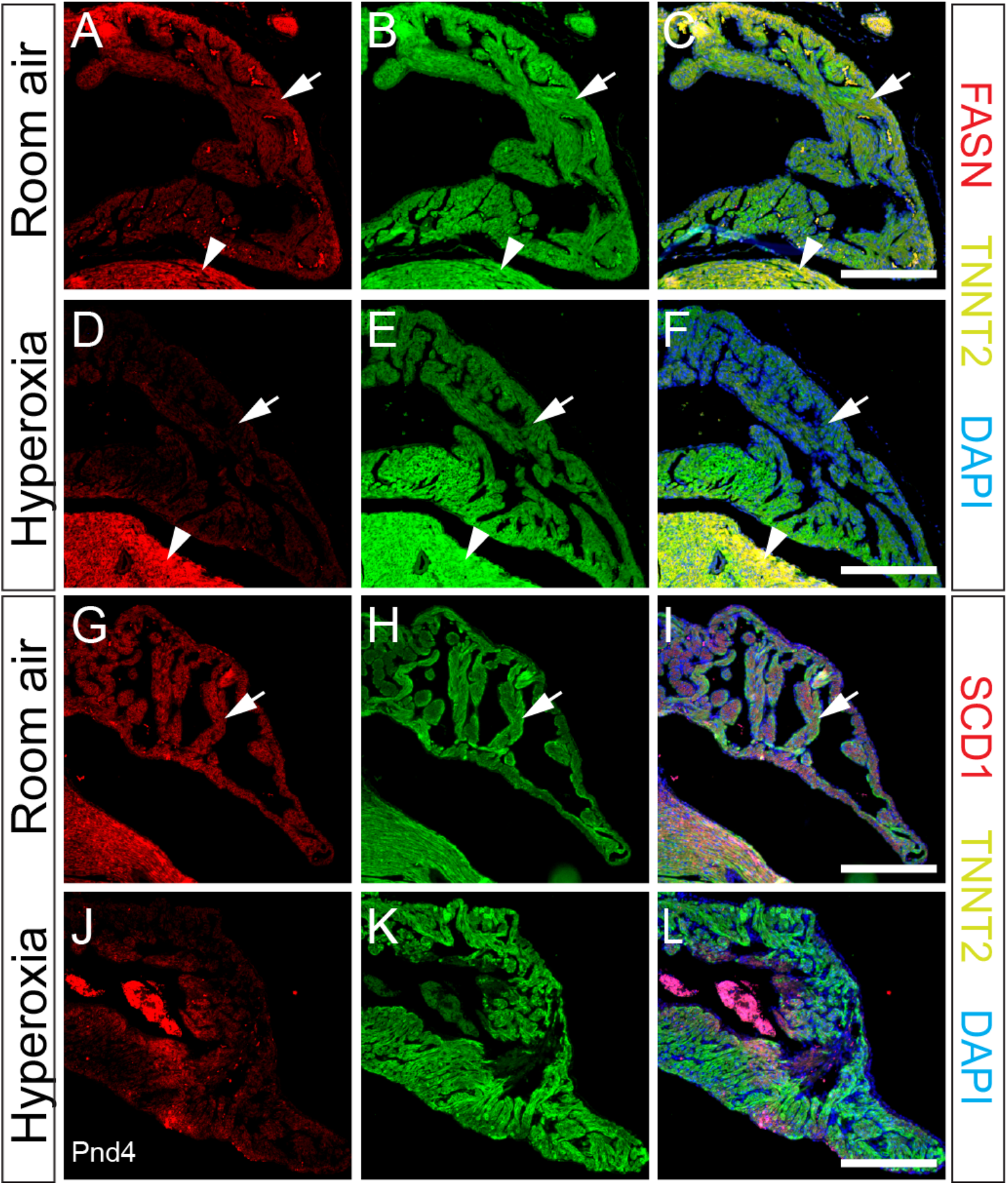
Neonatal hyperoxia represses fatty acid synthesis enzymes in the atrial cardiomyocytes of mice. (A-L) Sections through the left atrial appendages of PND4 neonates exposed to room air (A-C, G-I) or neonatal hyperoxia (D-F, J-L) co-stained for FASN (red, A, C, D, F) or SCD1 (red, G, I, J, L) and TNNT2 (green, B, C, E, F, H, I, K, L). Sections were also stained with DAPI to label nuclei (blue, C, F, I, L). Arrows and arrowheads show TNNT2+ cardiomyocytes in the left atrial appendage and LV, respectively. Scale bars = 200μm.

### Hyperoxia inhibits fatty acid synthesis genes and proliferation in the HL-1 line of immortalized murine atrial cardiomyocytes

To determine if the reduced expression of fatty acid synthesis genes in the atria of hyperoxia-exposed mice is a direct effect of oxygen on atrial cardiomyocytes, the levels of *Fasn* and *Scd1* mRNA were examined in HL-1 cells, a line of T-antigen immortalized atrial cardiomyocytes (42), that were exposed to room air (21% O_2_, 5% CO_2_) or hyperoxia (95% O_2_, 5% CO_2_) (**Figure 5A**). Relative to cells grown in room air, the expression of *Fasn* and *Scd1* were reduced in HL-1 cells grown in hyperoxia for 48 hours. Interestingly, while the levels of *Scd1* mRNA were lower in HL-1 cells grown in hyperoxia for 24 hours, *Fasn* was unaffected at this time. However, this difference in timing is unlikely to reflect a regulatory hierarchy between these genes because *Scd1* knockdown does not affect *Fasn* expression (data not shown). Together, these data suggest that hyperoxia can directly affect fatty acid synthesis genes in mouse atrial cardiomyocytes without the need for other cell types found within the heart.

**Figure 5.**
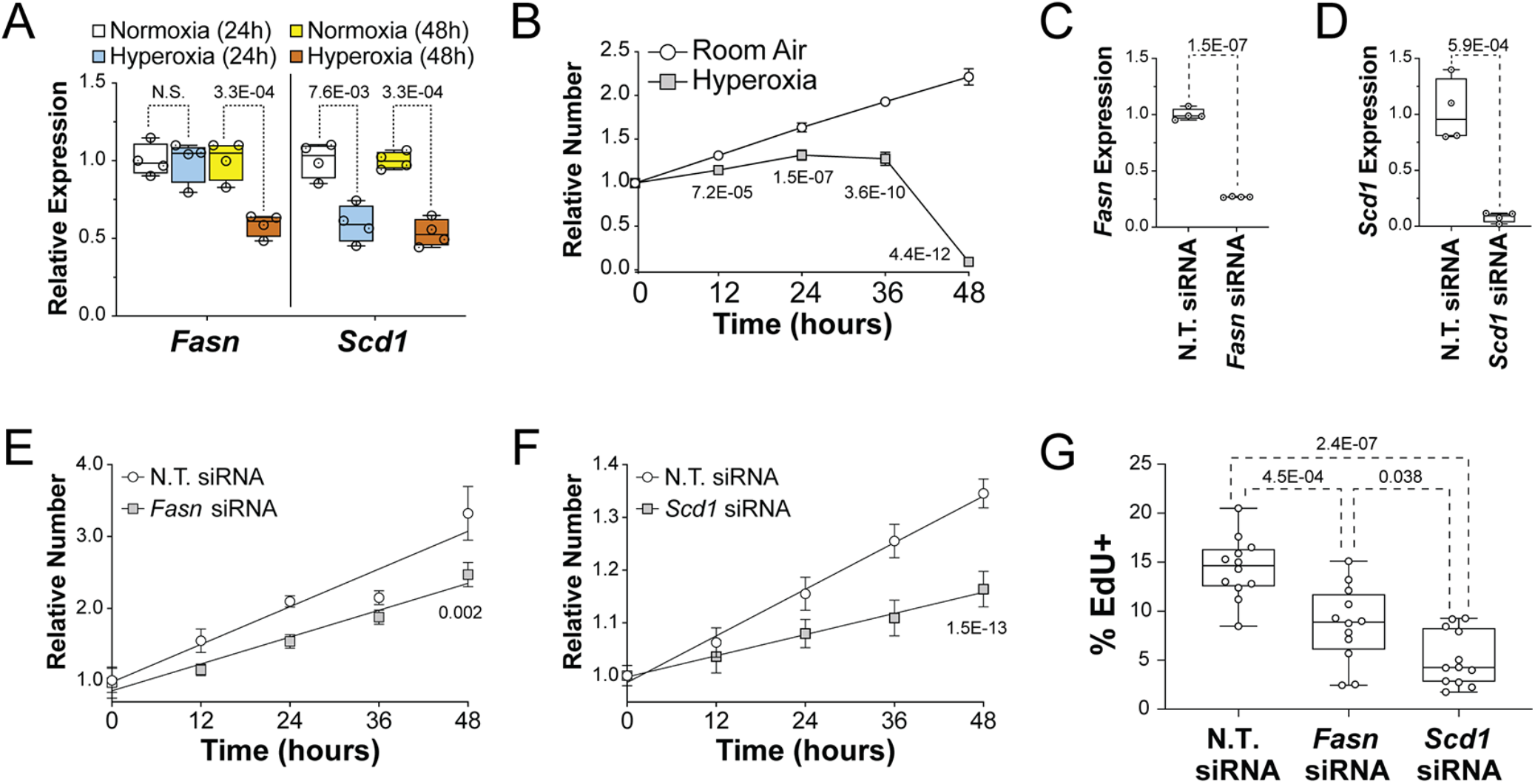
The suppression of fatty acid synthesis genes contributes to the reduced proliferation of HL-1 cells in hyperoxia. (A) Results of qRT-PCR for *Fasn* (left) and *Scd1* (right) in HL-1 cells grown in room air or hyperoxia for 24 hours (red and blue, respectively) or 48 hours (yellow and green, respectively). (B) Relative numbers of HL-1 cells plated at equal density and grown in room air (white circles) and hyperoxia (gray squares) for 48 hours relative to their density at 0 hours. (C and D) Results of qRT-PCR for *Fasn* (C) and *Scd1* (D) performed on HL-1 cells transfected *Fasn* and *Scd1* siRNAs, respectively. Control cells were transfected with non-targeting (N.T.) control siRNA. (D and E) Relative numbers of HL-1 cells transfected with control siRNA (white circles in E and F) and *Fasn* (Gray squares in E) or *Scd1* (Gray diamonds in F) siRNAs and grown for 48 hours in room air. Results are relative to the density of each group of cells at 0 hrs. (G) HL-1 cells were transfected with N.T., *Fasn* and *Scd1* siRNAs and grown in room air for 22 hours and treated with EdU for 2 hours before being fixed and stained. Graph shows the percentages of EdU labeled cells for each treatment. (B, E, and F) Error bars show 95% CI, lines and p-values are the results of linear regression analysis (A, C, D and G) Box plots show the median, 2^nd^ and 3^rd^ quartiles, error bars indicate range and markers show individual replicates. P-values are from one-way ANOVA with Holm-Sidak corrections (A and G) or unpaired t-tests (C and D).

To determine if the suppression of fatty synthesis genes in hyperoxia treated HL-1 cells correlates with reduced proliferation, asynchronously dividing HL-1 cells were plated at equal density and cultured in room air or hyperoxia for 0, 12, 24, 36 and 48 hours before being stained with DAPI and counted. HL-1 cells expanded continuously for 48-hours when grown in room air (**Figure 5B**). In contrast, HL-1 cells grown in hyperoxia expanded slower than controls for the first 24 hours, plateaued from 24 to 36 hours and then rapidly declined in numbers from 36 to 48 hours. These data indicate that hyperoxia directly represses proliferation and the expression of fatty acid synthesis genes in HL-1 atrial cardiomyocytes as it does in the atria of neonatal mice.

### Hyperoxia inhibits the proliferation of HL-1 atrial cardiomyocytes by repressing genes needed for fatty acid synthesis

To determine if fatty acid synthesis is required for atrial cardiomyocyte proliferation, HL-1 cells were transfected with non-targeting (N.T.) siRNA and siRNA for either *Fasn* or *Scd1* and grown in room air for 72 hours. Quantitative RT-PCR revealed that the levels of *Fasn* and *Scd1* mRNA were reduced around 75% and 90% in cells transfected with *Fasn* and *Scd1* siRNA, respectively, when compared to N.T. siRNA transfected controls (**Figure 5C, D**). Asynchronously dividing *Fasn, Scd1* and N.T. siRNA treated HL-1 cells were then plated at equal numbers and counted at 12-hour intervals (**Figure 5E, F**). While *Fasn, Scd1* and N.T. siRNA transfected cells expanded over the next 48 hours, cells transfected with *Fasn* and *Scd1* siRNA grew slower than N.T. siRNA transfected controls. To confirm that *Fasn* and *Scd1* are required for HL-1 cell proliferation, HL-1 cells were treated with DMSO as a vehicle control, the FASN inhibitor G28UCM at 10μM or the SCD1 inhibitor A939572 at 10nM (Tocris Biosciences) and grown in room air (**Supplemental Figure 1**). While cells in control media grew continuously for 72 hours, cells in A939572 plateaued after 24 hours and cells in G28UCM grew slower than controls during the first 24 hours and then declined in numbers from 24 and 72 hours. To more specifically examine the effects of *Fasn* and Scd1 knockdown on proliferation, *Fasn, Scd1* and N.T. siRNA transfected cells were grown for 23 hours, treated with EdU for 1 hour, fixed and stained with an anti-EdU antibody and DAPI. Scanning cells on an imaging cytometer showed that the fraction of cells with EdU incorporated into their DNA was reduced in *Fasn* and *Scd1* siRNA treated cells relative to N.T. siRNA transfected controls (**Figure 5G**). Taken together, these data indicate that reduced expression of *Fasn, Scd1* and other fatty acid synthesis genes may contribute to the reduced proliferation of atrial and pulmonary vein cardiomyocytes in mice exposed to neonatal hyperoxia.

### The overexpression of *Fasn* and *Scd1* restores the proliferation of HL-1 atrial cardiomyocytes in hyperoxia

To determine if overexpressing *Fasn* and *Scd1* would restore cardiomyocyte proliferation in hyperoxia, HL-1 cells transfected with empty vector or *Fasn* or *Scd1* expression vectors were plated at equal densities and grown in room air or hyperoxia for 0, 12 and 24 hours before being fixed, stained and counted. CMV-*Fasn* transfected cells expressed two fold more *Fasn* than cells transfected with empty vector (**Figure 6A**). As expected, control transfected cells grew slower in hyperoxia than in room air (**Figure 6B**). In contrast, the growth of CMV-*Fasn* transfected cells in room air and hyperoxia were equivalent to that of control cells in room air. Moreover, CMV-*Scd1* transfected cells expressed approximately 60 fold more *Scd1* than controls (**Figure 6C**) and grew faster than control cells in room air and hyperoxia (**Figure 6D**). To confirm that *Fasn* and *Scd1* expression restored HL-1 c proliferation in hyperoxia, cells were transfected with N.T., *Fasn* and *Scd1* plasmids, grown in room air or hyperoxia for 23 hours and treated with EdU to label cells in S-phase (**Figure 6E**). While *Fasn* and *Scd1* overexpression did not affect the numbers of EdU+ cells in room air, the fractions of *Fasn* and *Scd1* overexpressing cells that were EdU+ were higher than that for control cells in hyperoxia. *Fasn* and *Scd1* overexpression can thus partially restore the growth of HL-1 cell in hyperoxia.

**Figure 6.**
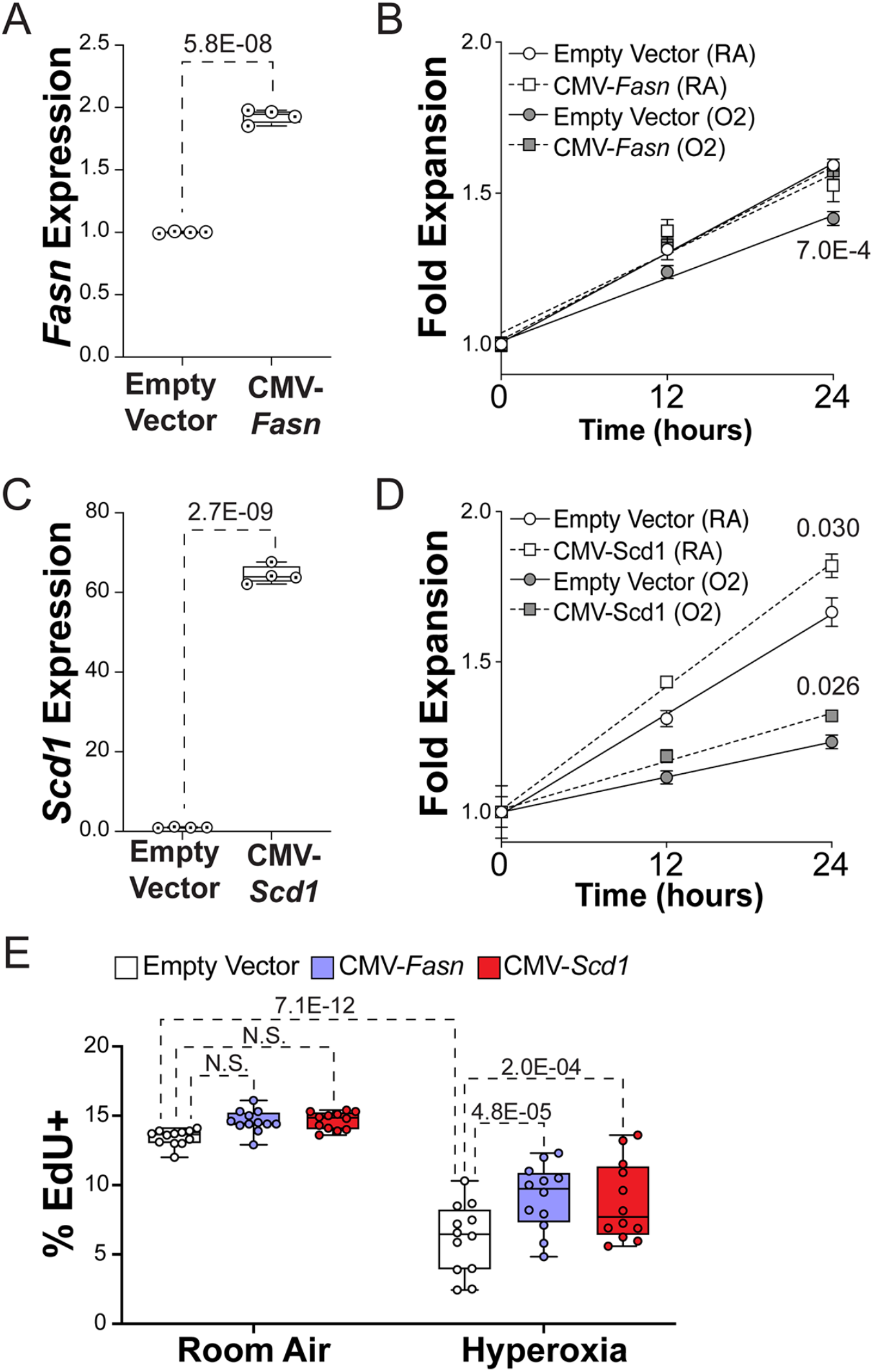
*Fasn* and *Scd1* overexpression increases the proliferation of HL-1 cells grown in hyperoxia. (A and C) Results of qRT-PCR for *Fasn* (A) and *Scd1* (C) in HL-1 cells 48 hours after they had been transfected with empty vector or *Fasn* (A) and *Scd1* (C) expression vectors. (B and D) The expansion of *Fas*n (B), *Scd1* (D) and control transfected HL-1 cells over 24 hours. Markers are the mean fold change in cell number for control (circles), *Fasn* (squares with dashed line, B) and *Scd1* (squares with dashed line, D) grown in room air (white) or hyperoxia (gray). Lines and p-values shown are the results of simple linear regressions. (E) Percentages of control (white), *Fasn* (blue) and *Scd1* (yellow) transfected HL-1 cells that incorporated EdU into their nuclei after 2 hours incubation in room air (left) and hyperoxia (right). (A, C, E) Box plots show the median, second and third quartiles, markers represent individual values, error bars show the range. P-values are the results of either unpaired t-tests (A and C) or one-way ANOVA with Holmes-Sidak corrections.

### Hyperoxia inhibits fatty acid synthesis genes and proliferation in human left atrial tissue explanted from infant donors

To determine if the changes observed in mice occur in humans, left atrial tissue from infants who died shortly after birth due to anencephaly was obtained from the Biorepository for Investigation of Neonatal Diseases of the Lung (BRINDL) at the University of Rochester. The muscle was separated from the surrounding tissue, cut into ~1 mm^3^ cubes, grouped and cultured in room air or hyperoxia for 24 hours before being lysed for RNA extraction and qRT-PCR or sectioned for immunologic staining. While hyperoxia did not alter the levels of *Fasn* mRNA, the expression of *Scd1* was reduced in hyperoxia-exposed explants relative to controls (**Figure 7A**). Immunological staining showed that SCD1 protein localized to TNNT2+ cardiomyocytes in explants exposed to room air but not in those exposed to hyperoxia (**arrows, Figure 7B**). Sectioned explants were stained for the proliferation marker Ki-67 and TNNT2 to determine if hyperoxia reduced the proliferation of human left atrial cardiomyocytes. Ki67+ nuclei were more frequently observed in TNNT2+ cardiomyocytes exposed to room air than in those exposed to hyperoxia (**arrows, Figure 7C**), more these cells were found in explants exposed to room air (**Figure 7D**). Together, these data indicate that hyperoxia suppresses fatty acid synthesis genes and proliferation in human left atrial tissue as it does in mice.

**Figure 7.**
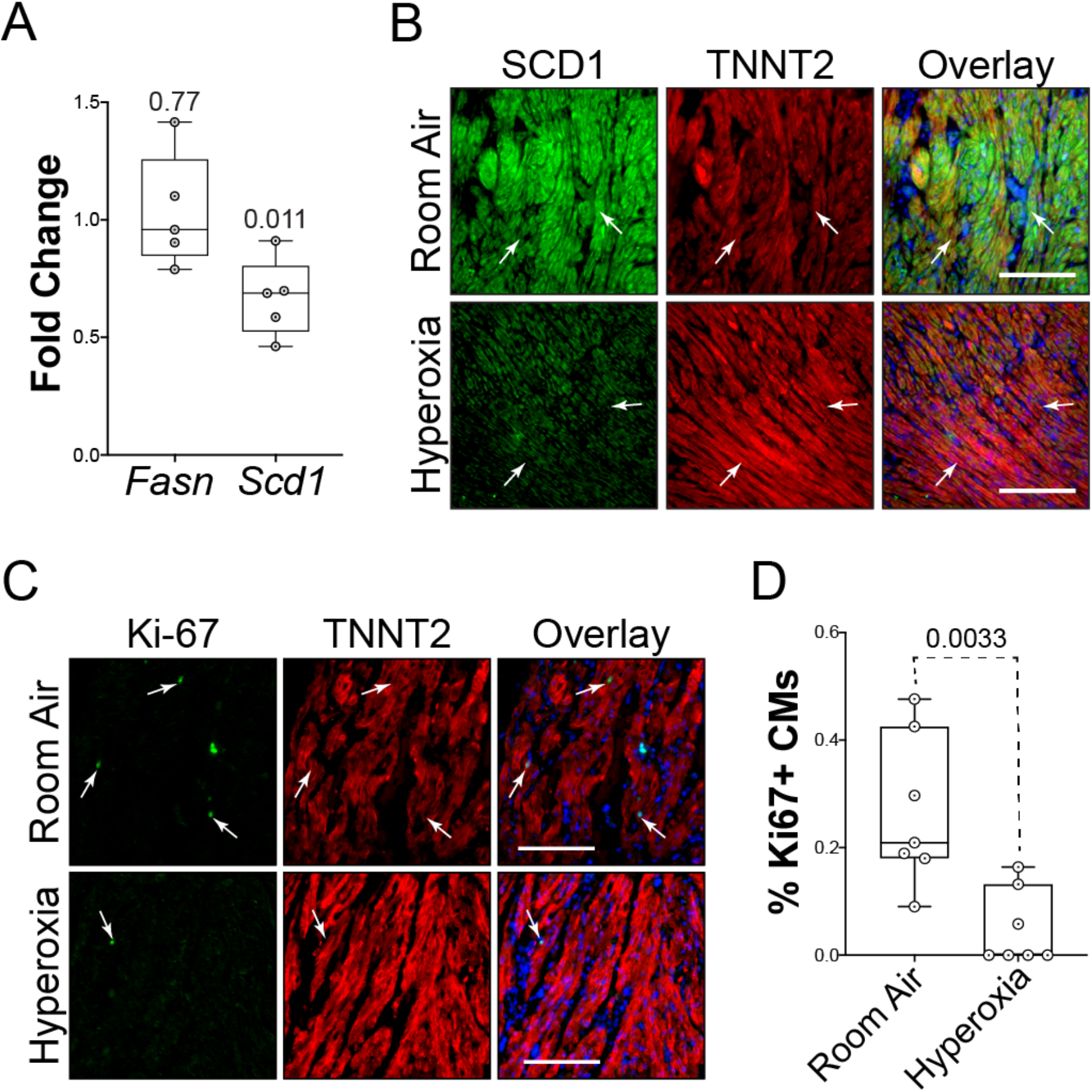
Hyperoxia suppresses fatty acid synthesis genes in left atrial tissue explanted from infant human donors. (A) Results of qRT-PCR for *Fasn* and *Scd1* in left atrial tissue explanted from 5 human infants who died at birth due to anencephaly and exposed to room air or hyperoxia for 24 hours. (B) Sections of explants exposed to room air (top) or hyperoxia (bottom) were stained for SCD1 (green), TNNT2 (red) and DAPI (blue). (C) Sections of explants exposed to room air (top) and hyperoxia (bottom) stained for the proliferation marker Ki67 (green), TNNT2 (red) and DAPI (blue). (D) Graph shows the percentages of TNNT2 expressing cells with Ki67 positive nuclei in explants exposed to room air or hyperoxia. (A and D) Circles indicate fold changes in gene expression between hyperoxia-exposed and control explants from 7 individual donors (A) or percentages of TNNT2 labeled cells with Ki67 positive nuclei in sections (D). Boxes are the median, 2^nd^ and 3^rd^ quartiles, error bars show range and p-values are the results of either single sample t and Wilcox tests (A) or unpaired t-tests (D). Scale bars = 100μm.

## DISCUSSION

There is a significant need to understand why high oxygen exposure at birth is a risk factor for adult cardiovascular disease. We previously showed that neonatal hyperoxia causes pulmonary hypertension in adult mice and that this phenotype was preceded by a loss of the cardiomyocytes surrounding the pulmonary vein due to reduced proliferation (29). We now extend these findings by showing that hyperoxia similarly inhibits the postnatal proliferation of left atrial cardiomyocytes and causes the myocardium of this chamber to be hypoplastic. While the remaining cardiomyocytes become hypertrophic to compensate for their reduced numbers, the left atria of hyperoxia-exposed mice dilate later in life and lose their ability to pump blood into the left ventricle during diastole. The loss of left ventricular filling reduced the diastolic volume, stroke volume and cardiac output of this chamber in aged mice exposed to neonatal hyperoxia without affecting left ventricular systolic function. Neonatal hyperoxia may thus initiate the development of adult diastolic dysfunction by disrupting the postnatal proliferation of left atrial cardiomyocytes.

Affymetrix arrays, qRT-PCR and immunohistochemistry revealed that neonatal hyperoxia inhibited genes needed for *de novo* fatty acid synthesis in the atrial but not ventricular cardiomyocytes of mice exposed to neonatal hyperoxia. These included *Fasn* and *Scd1,* which are elevated in highly proliferative tumors, increase proliferation when overexpressed and required for the proliferation of non-cardiac cells (43–49). We thus explored whether the persistent repression of *Fasn* and *Scd1* expression is responsible for the reduced proliferation of left atrial cardiomyocytes in mice exposed to neonatal hyperoxia. Left atrial cardiomyocytes can be isolated from neonatal mice but the yield is low and the resulting cells lose their proliferative capacity quickly in culture, making it difficult to examine how genetic manipulations affect their expansion. The HL-1 line of immortalized atrial cardiomyocytes was thus chosen to model the effects of hyperoxia on neonatal atrial cardiomyocytes since these cells are differentiated and contractile but still metabolically immature and proliferative (42, 50, 51). Consistent with the loss of atrial CM proliferation in hyperoxia-exposed mice, exposing HL-1 cells to hyperoxia for 48 hours reduced *Fasn* and *Scd1* levels and proliferation, confirming that hyperoxia can directly affect atrial cardiomyocytes in the absence of other cell types within the heart. Chemical inhibitors and siRNAs for *Fasn* and *Scd1* also reduced HL-1 cell proliferation while overexpressing these genes partially restored their expansion in hyperoxia, further suggesting repression of *Fasn, Scd1* and other fatty acid synthesis genes mediate the inhibitory effects of neonatal hyperoxia on left atrial cardiomyocyte proliferation (52). These data suggest that neonatal hyperoxia inhibits left atrial cardiomyocyte proliferation by reducing the expression of Fasn, Scd1 and other fatty acid synthesis genes within these cells.

A prior study found hyperoxia inhibits proliferation of ventricular cardiomyocyte by activating ATM-dependent DNA damage signaling (53), however, ATM target genes were unaffected in our microarray study of atria from hyperoxia-exposed and control mice. *Fasn, Scd1* and other fatty acid synthesis genes were also unaffected in the ventricles of the mice exposed to neonatal hyperoxia, suggesting hyperoxia affects the proliferation of atrial and ventricular cardiomyocytes through distinct mechanisms. Since the left atrium is the first chamber to encounter oxygen-rich blood from the lungs, left atrial cardiomyocytes may have evolved to have a different response to hyperoxia than ventricular cardiomyocytes. Activation of ATM signaling in these cells could also reflect an alternative pathway that is activated when the mice were exposed to hyperoxia during late embryogenesis, a time when Fasn expression is normally declining in ventricular cardiomyocytes (54). This ATM-dependent suppression of ventricular cardiomyocyte proliferation was hypothesized to reflect a role for oxygen in promoting the terminal differentiation of these cells. However, in our model, *Fasn, Scd1* and other fatty acid synthesis genes continue to be suppressed in the atria of mice exposed to hyperoxia even after they returned to room air. If neonatal hyperoxia accelerated the normal differentiation of atrial cardiomyocytes as previously proposed, fatty acid synthesis genes should return to similar levels in the atria of hyperoxia-exposed and control mice as the atrial cardiomyocytes of control mice matured during development. These data thus indicate that exposure to hyperoxia in early neonatal life permanently reprograms atrial cardiomyocytes metabolism in a maladaptive fashion that contributes to the long-term effects of neonatal hyperoxia on cardiovascular health. Whether persist molecular changes exist in ventricular cardiomyocytes remains to be determined.

The roles for fatty acid synthesis in cellular proliferation are not fully understood. Oxidative phosphorylation in Melan-A cells, a line of immortalized melanocytes can be suppressed with pharmacologic inhibitors or siRNA to FASN (44). This loss of mitochondrial function was partially due to the accumulation of malonyl-CoA, which inhibits the transports fatty acids into mitochondria at high concentrations. FASN inhibition can also reduce mitochondrial inner membrane potential (ΔΨm) and increase the production of mitochondrial reactive oxygen species (55, 56). As observed with FASN, pharmacological and genetic deletion of SCD1 activity reduces proliferation and survival of cells disrupts mitochondrial function, increases ROS production and induces apoptosis in many cell types including ventricular cardiomyocytes isolated from neonatal mice (52, 57). Since SCD1 converts saturated fatty acids to the mono-unsaturated forms used to make phospholipids, modulating SCD1 function may affect the availability of membranes for newly replicated cells. SCD1 inhibition also reduces mitochondrial function by disrupting their membrane composition and increasing the leakage of protons. The loss of SCD1 function also activates AMP-dependent protein kinase (AMPK) in many contexts, including the hearts of Scd1 knockout mice (36, 58, 59). AMPK phosphorylates and inhibits the pro-proliferative serine/ threonine kinase mammalian target of rapamycin (mTOR) (60), and Serum response element binding protein (SREBP), the master regulator of fatty acid synthesis gene transcription (61).

Recent echocardiographic, magnetic resonance and CT imaging studies indicate that preterm infants have larger left atria and lower diastolic function than term infants (14–17), suggesting that the heart failure among young adults who were born preterm has developmental origins. Serum from preterm infants was shown to have higher levels of malonyl-carnitine and lower levels of palmatoyl-carnitine than serum from term infants born (62), suggesting a link between preterm birth and reduced FASN activity. Preterm infants often require mechanical ventilation, steroids, and other therapeutic interventions that make it difficult to identify the direct effects of supplemental oxygen on their cardiovascular health. We thus sought to determine if neonatal hyperoxia suppresses fatty acid synthesis genes and proliferation in explanted human atrial cardiac slices. Myocardial tissue was thus explanted from the left atria of human infant donors who died at birth due to anencephaly and cultured in room air or hyperoxia for 24 hours before *Fasn* and *Scd1* were assessed by qRT-PCR and immunological staining. Sectioned explants were also stained for Ki67 to determine if hyperoxia affected cardiomyocyte proliferation in humans as it does in mice. Hyperoxia reduced *Scd1* expression and the numbers of Ki67+ cardiomyocytes in explanted atrial tissue, confirming hyperoxia represses at least one major component of the fatty acid synthesis pathway and proliferation in the left atrial cardiomyocytes of human infants. While *Fasn* was unaffected in explanted human atrial tissue exposed to hyperoxia for 24 hours, one day of exposure was similarly insufficient to repress *Fasn* in HL-1 cells. Prolonged exposure may thus be required for hyperoxia to affect *Fasn* expression in human atrial cardiomyocytes as it does in the mice. We acknowledge these studies used tissue from anencephalic infants that were born at term and that oxygen sensitivity is likely to change over the course of gestation and that these *ex vivo* studies may not fully recapitulate the effects of neonatal hyperoxia observed *in vivo*. Despite these limitations, our findings suggest the response of left atrial tissue from newborn human to hyperoxia parallels that of mice and HL-1 cells and that altered fatty acid metabolism may underlie the cardiovascular disease in preterm infants.

The data herein suggest that exposure to hyperoxia in early postnatal life initiates the development of adult diastolic dysfunction by permanently suppressing *Fasn, Scd1* and other genes needed for fatty acid synthesis and cardiomyocyte proliferation in the walls of the left atrium and pulmonary vein. Moreover, since adult cardiomyocytes rely on fatty acids for 70% of the ATP used for contractility, the long-term suppression of *Fasn, Scd1* and other fatty acid synthesis genes may also affect the functionality of the remaining left atrial cardiomyocytes after they have stopped proliferating and further contribute to the diastolic heart failure that follows neonatal hyperoxia. Pharmacological interventions that promote fatty acid synthesis may thus prove useful for dampening the long-term deleterious effects of neonatal hyperoxia on the cardiovascular health of former preterm infants.

## MATERIALS AND METHODS

### Mice and hyperoxia

C57BL/6J mice were purchased from The Jackson Laboratories (Bar Harbor, MA) and maintained as an inbred colony. The mice were exposed to humidified room air (21% oxygen) or hyperoxia (100% oxygen) between postnatal days 0-4 (22). Some mice exposed to hyperoxia were then returned to room air. Dams were cycled between room air and hyperoxia every 24 hours to reduce oxidant injury to their lungs. The mice were provided food and water *ad libitum* and housed in micro-isolator cages in a specified pathogen-free environment according to a protocol approved by the University Committee on Animal Resources at the University of Rochester.

### Echocardiography

To assess cardiac function, mice were anesthetized with isoflurane and subject to transthoracic echocardiography using a VisualSonics Vevo 3100. Readings were obtained with a 40 MHz transducer in M-mode along the short axis of the LV. Measurements were made by the staff of the Microsurgery and Echocardiography Core at the Aab Cardiovascular Research Institute, who were blinded to the genotype of the animals.

### Affymetrix Arrays and Analysis

Total RNA was isolated from the atria of PND4 mice exposed to room air or hyperoxia using Trizol (ThermoFisher Scientific, Waltham, MA) and its integrity was validated using an Agilent Bioanalyzer (Agilent Technologies, Santa Clara, CA). The RNA was converted to cDNA, biotinylated with Ovation kits from NuGEN (San Carlos, CA) and hybridized to the Affymetrix mouse genome 430 2.0 array (Affymetrix, Santa Clara, CA). Each array was probed with RNA isolated from an individual mouse. Arrays were stained with streptavidin-phycoerythrin as recommended by Affymetrix. The arrays were then scanned for phycoerythrin fluorescence and the spot intensities were normalized across arrays with the RMA method in the “oligo” package (63). Affymetric cell file were loaded in R 3.4.3/Bioconductor 3.5 using the oligo package and RMA-normalized. 29290 features that passed a filters on mean expression > 3 and total variance > 0.0025 were tested for differential expression with limma (64). A total of 157 genes with FDR-adjusted q-values < 0.30 was tested for enrichment in the gene ontology using clusterProfiler. The complete array dataset was deposited in ArrayExpress under accession number E-MTAB-9008.

### Quantitative RT-PCR

Total RNA was isolated using Trizol reagent, treated with DNase 1 to remove genomic DNA and reverse transcribed using the Maxima First Strand cDNA synthesis kit (ThermoFisher Scientific, Waltham, MA). Quantitative reverse transcriptase polymerase chain reaction (qRT-PCR) was performed using the primer pairs listed in **Table 4** and iTaq Universal SYBR Green Master Mix on a CFX384™ Real-Time PCR detection system (Bio-Rad Laboratories, Hercules, CA). Fold changes in gene expression were calculated by the ΔΔCt method using the weighted average of Ct values for the housekeeping genes *Pol2ra, Tbp* and *Gapdh* to control for loading.

### Tissue processing, histological sectioning and Immunostaining

Intraperitoneal injections of Avertin and heparin were used to euthanize mice and prevent clotting, respectively. Hearts were perfused with PBS and 10% neutral buffered formalin (NBF) to remove blood cells and fix the tissue, respectively and fixed in 4% PFA and fixed overnight before being embedded in paraffin, sectioned, and stained as described (20, 65). Sections were either stained with hematoxylin and eosin (H&E) and Masson’s trichrome to assess morphology or antibodies for FASN and SCD1 (Thermo-Fisher, PA5-19509 and PA5-17409, respectively) and fluorescently labeled secondary antibody (Jackson Immune Research, West Grove, PA) to examine protein levels and localizations. Co-staining with an antibody for TNNT2 (Thermo-Fisher, MA5-12960) and 4’, 6-diamidino-2-phenylindole (DAPI) were used to label cardiomyocytes and nuclei, respectively. Sections were imaged using a Nikon E800 Fluorescence microscope (Microvideo Instruments, Avon, MA) and a SPOT-RT digital camera (Diagnostic Instruments, Sterling Heights, MI).

### Culture of immortalized murine atrial cardiomyocytes

HL-1 cells were cultured in Claycomb media with 100 μM norepinephrine, 300 μM ascorbic acid, 10% fetal bovine serum, 4 mM glutamine and penicillin/ streptomycin (Sigma-Aldrich, St. Louis, MO). When indicated, cells were transfected with pools of siRNA against *Fasn* and *Scd1* (Horizon, L-040091-00 and M-040675-01) or *Fasn* and *Scd1* cDNAs (Horizon, MMM1013-202765185 and MMM1013-202762947) with Lipofectamine RNAiMax or 2000 (Thermo-Fisher, Waltham, MA). Cells transfected with non-targeting siRNA and empty vector served as controls. For growth assays, cells were plated in 96 well dishes at a density of 5000 viable cells/ well and allowed to attach overnight. Cells were then exposed to room air and hyperoxia for 48 hours with plates being fixed at 0, 12, 24 and 48 hours. Fixed cells were stained with Hoechst and scanned on a Celigo S Image Cytometer (Nexcelom Bioscience, Lawrence, MA) to count the nuclei in each well using the associated software. When indicated, 10 mM G28UCM and 10 nM A939657 (Tocris Bioscience, Bristol, UK) were added prior to exposure to inhibit FASN and SCD1, respectively. Cells treated with DMSO were used as vehicle only controls. EdU incorporation, cells were exposed to room air or hyperoxia for 24 hours before 10 μM 5-ethynyl-2’-deoxyuridine (EdU) was added to the media. Cells were then returned to room air or hyperoxia for 1 hour and fixed before EdU+ cells were detected with the Click-IT EdU Cell Proliferation Kit (Thermo-Fisher, Waltham, MA). Cells were costained with Hoechst and imaged on a Celigo S Image Cytometer to determine the percentages of cells in each well with EdU incorporated into their nuclei.

### Ex vivo culture of human left atrial tissue

Left atrial tissue samples were provided through the federal United Network of Organ Sharing via the International Institute for Advancement of Medicine (IIAM) and entered into the NHLBI LungMAP Biorepository for INvestigations of Diseases of the Lung (BRINDL) at the University of Rochester Medical Center overseen by the IRB as RSRB00047606, as previously described (66, 67). Tissue from 4 donors with anencephaly who died within 24 hours of birth at 37 weeks and 1 donor with Hirschsprung’s and demyelinating disease that died at 3 months of age was used to examine the effects of hyperoxia on *Fasn, Scd1* and *Ki67* expression. Explants from these donors and an additional 2 donors near birth due to anencephaly for which RNA was unavailable were sectioned and used to examine the effects of hyperoxia on staining for SCD1 and Ki67 protein. In all cases, the left atrial myocardium was separated from the surrounding tissue, cut into ~1 mm^3^ cubes and cultured in serum free EMEM supplemented with Insulin-Transferrin-Selenium (ITS). Explants were examined on an inverted microscope with a heated stage to confirm that they were beating and viable before being divided into groups of 10-15 explants and cultured in room air or hyperoxia for 24 hours as described. After exposure, explants were again examined on an inverted microscope to confirm viability and then either lysed for RNA extraction and qRT-PCR or fixed for histological sectioning and immunologic staining. For qRT-PCR studies, the fold change in gene expression between explants cultured in room air and hyperoxia were calculated for five donors and averaged to report mean fold changes in mRNA levels. One sample t-tests were used to determine if mean fold changes relative to room air exposed explants in *Fasn* and *Scd1* deviated significantly from one.

### Statistical analysis

Data was analyzed with JMP12 software (SAS Institute, Cary, NC) and graphed with Prism 8 (GraphPad Software, San Diego, CA). Differences in the left atrial area, cellular density, proliferation and other single variant studies were judged using unpaired t-tests. Differences in cardiac function, gene expression and other simultaneously measured parameters were judged with unpaired t-tests and Holm-Sidak corrections. Changes in EdU labeling and other studies with more than three experimental conditions were judged using one-way ANOVA with Tukey’s multiple comparisons tests. Differences in HL-1 cell expansion were judged by two-way ANOVA with Sidak multiple comparisons tests or by linear regression analysis. In all cases, p < 0.05 were considered to be significant. The statistical analysis of Affymetrix array data and gene expression in explanted human atrial tissue was performed as described in their respective subsections of the Materials and Methods.

## ACKNOWLEDGEMENTS

This work was funded in part by National Institutes of Health Grants R01 HL091968 (M.A. O’Reilly), Strong Children’s Research Center pilot grant (E.D. Cohen), and a Wine Auction Pilot from the Cardiovascular Research Institute (M.A. O’Reilly and G. Porter). NIH Center Grant P30 ES001247 supported the animal inhalation facility and tissue-processing core. The University of Rochester’s Department of Pediatrics provided financial support through the Perinatal and Pediatric Origins of Disease Program and directly to the Pediatric Histology Service. We thank Robert Gelein for maintaining the oxygen exposure facility, Daria Krenitsky for tissue processing and sectioning, and the University of Rochester’s Genomic Research Center for sequencing mRNA. We thank the donating families, the Research Recovery Organization IIAM, the LungMAP Consortium supported by NHLBI Molecular Atlas of Lung Development Program Human Tissue Core grants (U01HL122700 and U01HL148861) and the members of the Pryhuber laboratory, Heidie Huyck and Cory Poole, who assisted with preparation of the human heart tissue.

## SUPPLEMENTAL DATA SECTION

**Supplemental Figure 1.**
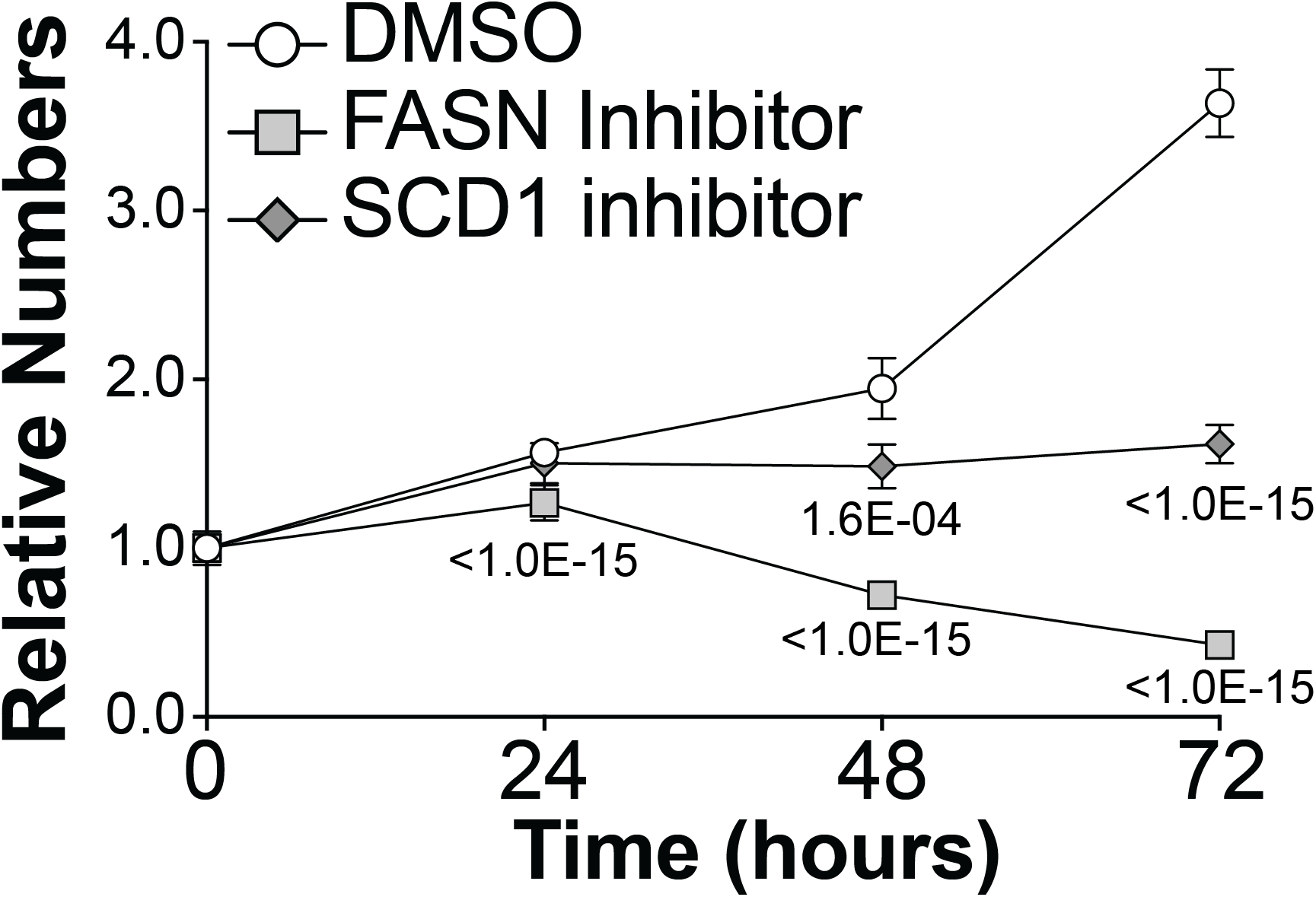
Effects of FASN and SCD1 inhibitors on HL-1 cell growth. Graph shows the increase in the relative numbers of HL-1 cells grown for 72 hours in media containing DMSO (white circles), the FASN inhibitor G28UCM at 10 mM (gray squares), or the SCD1 inhibitor A939572 at 10 nM (gray diamonds). Values are normalized to number of cells in each well just prior to adding inhibitor (0hrs). The p-values shown are the results of two-way ANOVA with Dunnett’s multiple comparison tests.

**Supplemental Table 1.** Results of echocardiographs performed on Pnd56 and Pnd365 mice exposed to room air or hyperoxia at birth.

| Attribute | Pnd56 |  |  | Pnd365 |  |  |
| --- | --- | --- | --- | --- | --- | --- |
|  | Room Air | Hyperoxia | *p*-value | Room Air | Hyperoxia | *p*-value |
| Heart Rate (BPM) | 455.06 ± 61.59 | 539.43 ± 62.54 | 0.06384 | 568.91 ± 36.05 | 556.06 ± 31.08 | 0.56274 |
| Diameter; systole (mm) | 2.76 ± 0.42 | 2.51 ± 0.31 | 0.30323 | 1.82 ± 0.14 | 1.80 ± 0.14 | 0.81281 |
| Diameter; diastole (mm) | 4.00 ± 0.38 | 3.71 ± 0.19 | 0.16940 | 3.40 ± 0.13 | 3.03 ± 0.19 | 0.00739* |
| Volume; systole (μL) | 29.49 ± 11.30 | 22.94 ± 7.38 | 0.30930 | 10.17 ± 1.95 | 9.85 ± 1.86 | 0.79556 |
| Volume; diastole (μL) | 70.64 ± 16.08 | 58.76 ± 7.25 | 0.17063 | 47.60 ± 4.15 | 36.17 ± 5.78 | 0.00705* |
| Stroke Volume (μL) | 41.15 ± 4.96 | 35.82 ± 2.50 | 0.06458 | 37.43 ± 2.98 | 26.32 ± 5.25 | 0.00339* |
| Ejection Fraction (%) | 59.34 ± 6.17 | 61.61 ± 7.38 | 0.61266 | 78.73 ± 2.98 | 72.53 ± 5.27 | 0.05107* |
| Fractional Shortening (%) | 31.18 ± 3.99 | 32.66 ± 4.85 | 0.61424 | 46.43 ± 2.90 | 40.52 ± 4.48 | 0.03824* |
| Cardiac Output (mL/min) | 18.49 ± 0.75 | 19.37 ± 2.93 | 0.53534 | 21.31 ± 2.54 | 14.59 ± 2.59 | 0.00324* |
| LV Mass (mg) | 106.97 ± 14.45 | 87.33 ± 9.64 | 0.03534* | 74.42 ± 5.18 | 67.33 ± 14.81 | 0.34218 |
| LV Mass Cor (mg) | 85.57 ± 11.56 | 69.86 ± 7.71 | 0.03534* | 59.53 ± 4.14 | 53.87 ± 11.85 | 0.34218 |
| LV Anterior Wall Diameter, Systole (mm) | 1.22 ± 0.17 | 1.16 ± 0.08 | 0.44820 | 1.20 ± 0.05 | 1.17 ± 0.11 | 0.51302 |
| LV Anterior Wall Diameter, Diastole (mm) | 0.80 ± 0.09 | 0.74 ± 0.08 | 0.24519 | 0.76 ± 0.06 | 0.76 ± 0.06 | 0.88400 |
| LV Posterior Wall Diameter, Systole (mm) | 1.06 ± 0.26 | 1.02 ± 0.18 | 0.79105 | 1.03 ± 0.05 | 1.01 ± 0.09 | 0.71258 |
| LV Posterior Wall Diameter, Diastole (mm) | 0.69 ± 0.17 | 0.67 ± 0.14 | 0.82254 | 0.63 ± 0.06 | 0.71 ± 0.14 | 0.28208 |

Cardiac function measurements of young (Pnd56) and old (Pnd365) mice exposed to room air or hyperoxia as neonates. Values represents mean ± standard deviation of 5 mice per group. \* $p \leq 0.05$ relative to room air.

**Supplemental Table 2.**
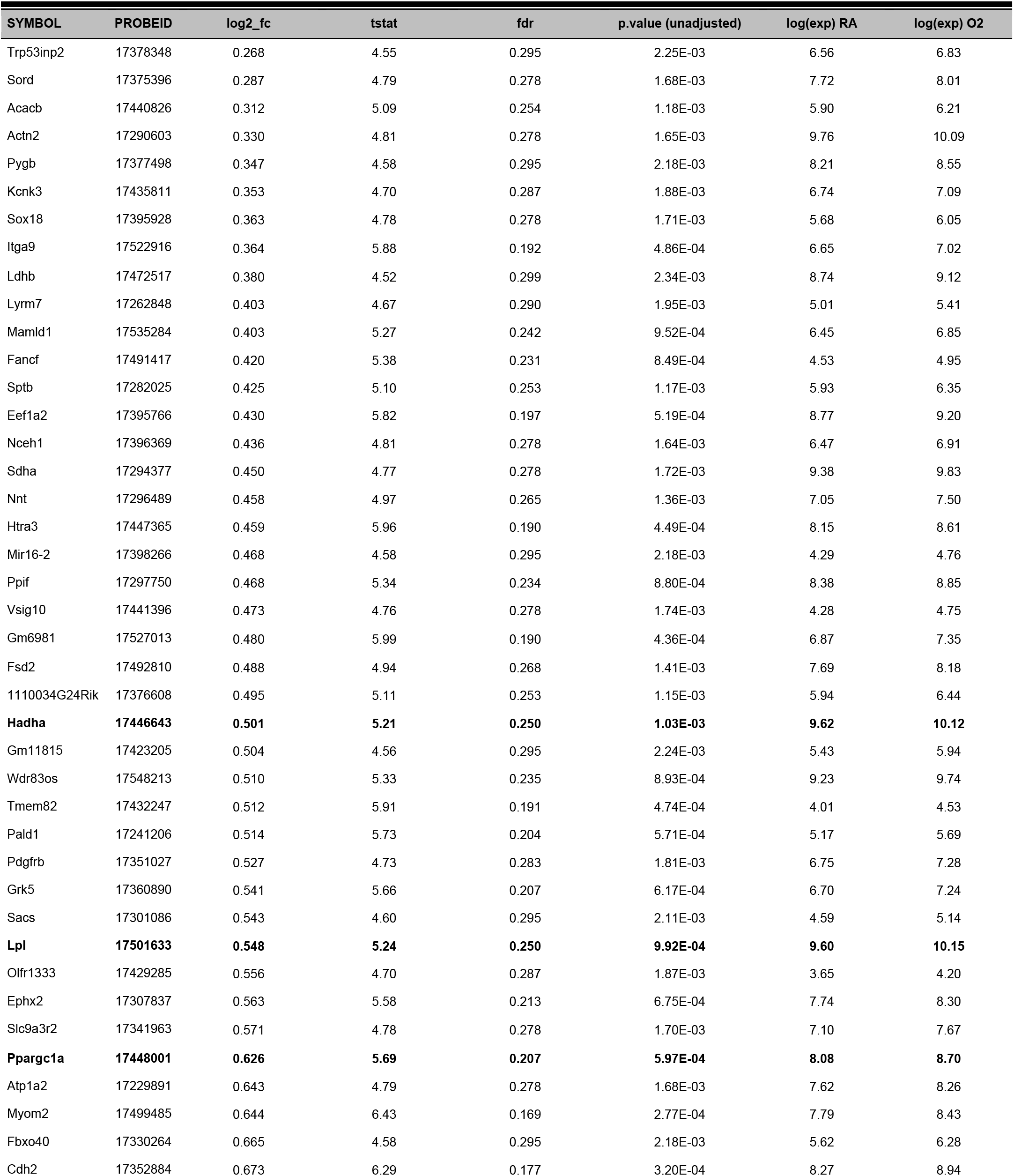
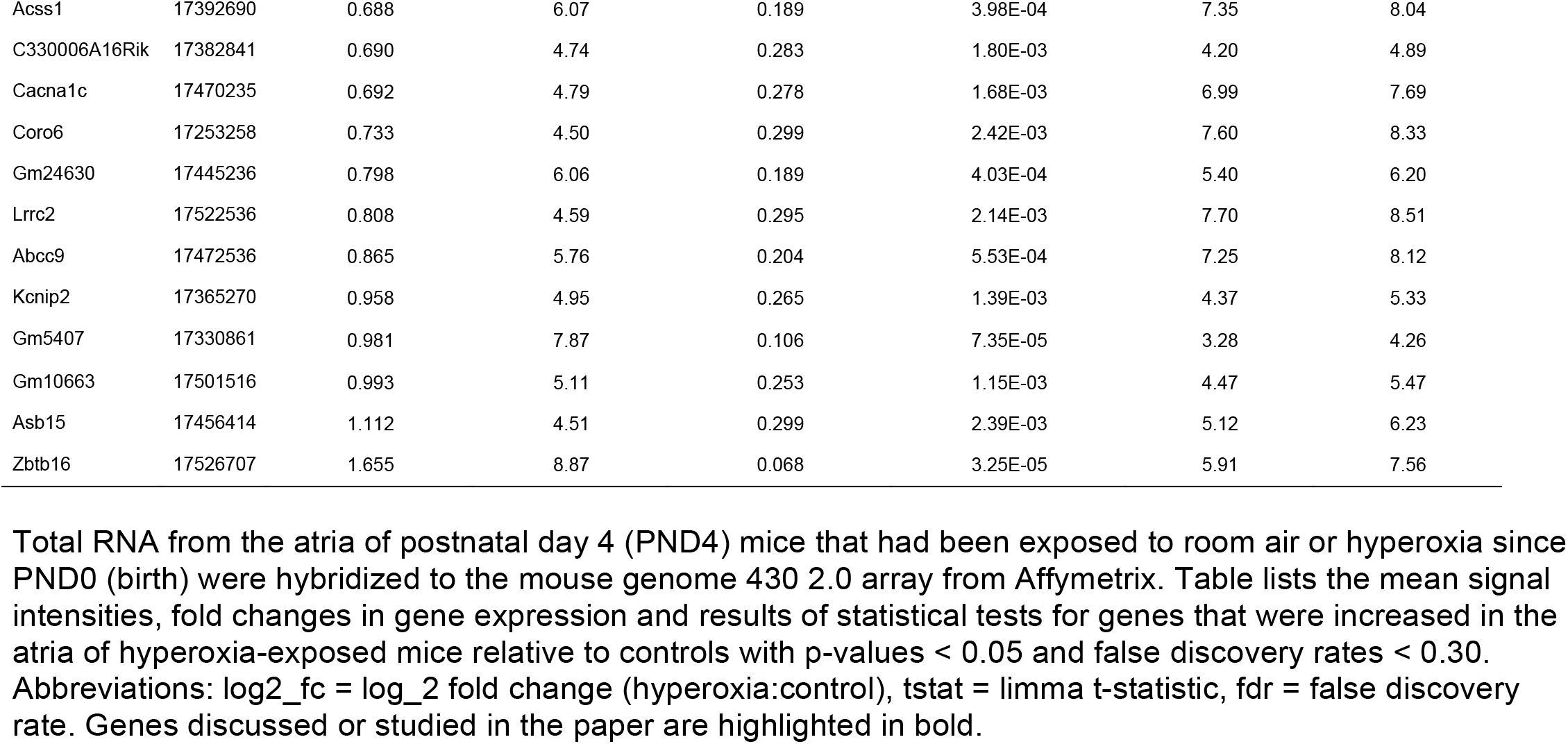
Genes expressed at higher levels in the atria of neonatal hyperoxia-exposed mice relative to controls.

**Supplemental Table 3.**
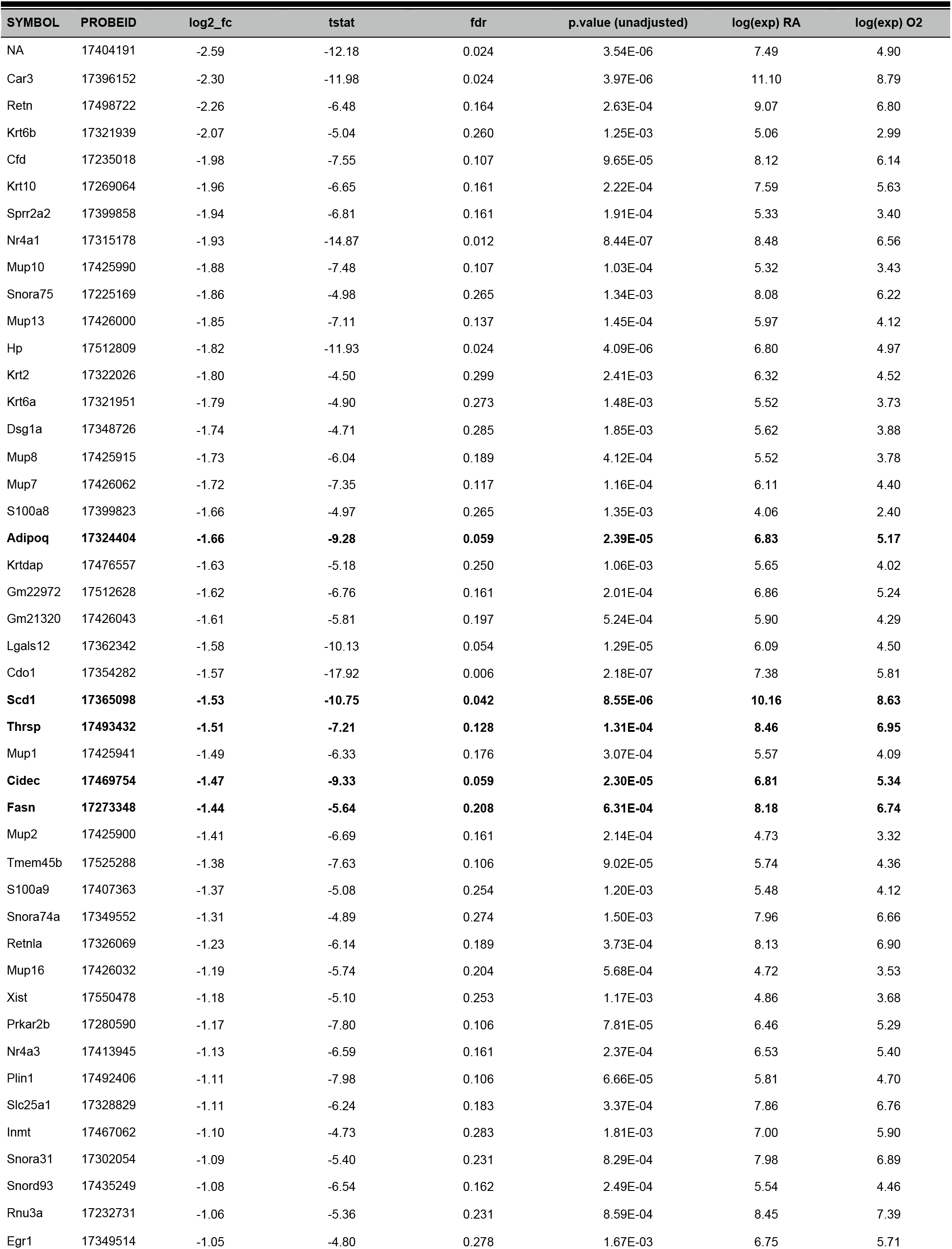
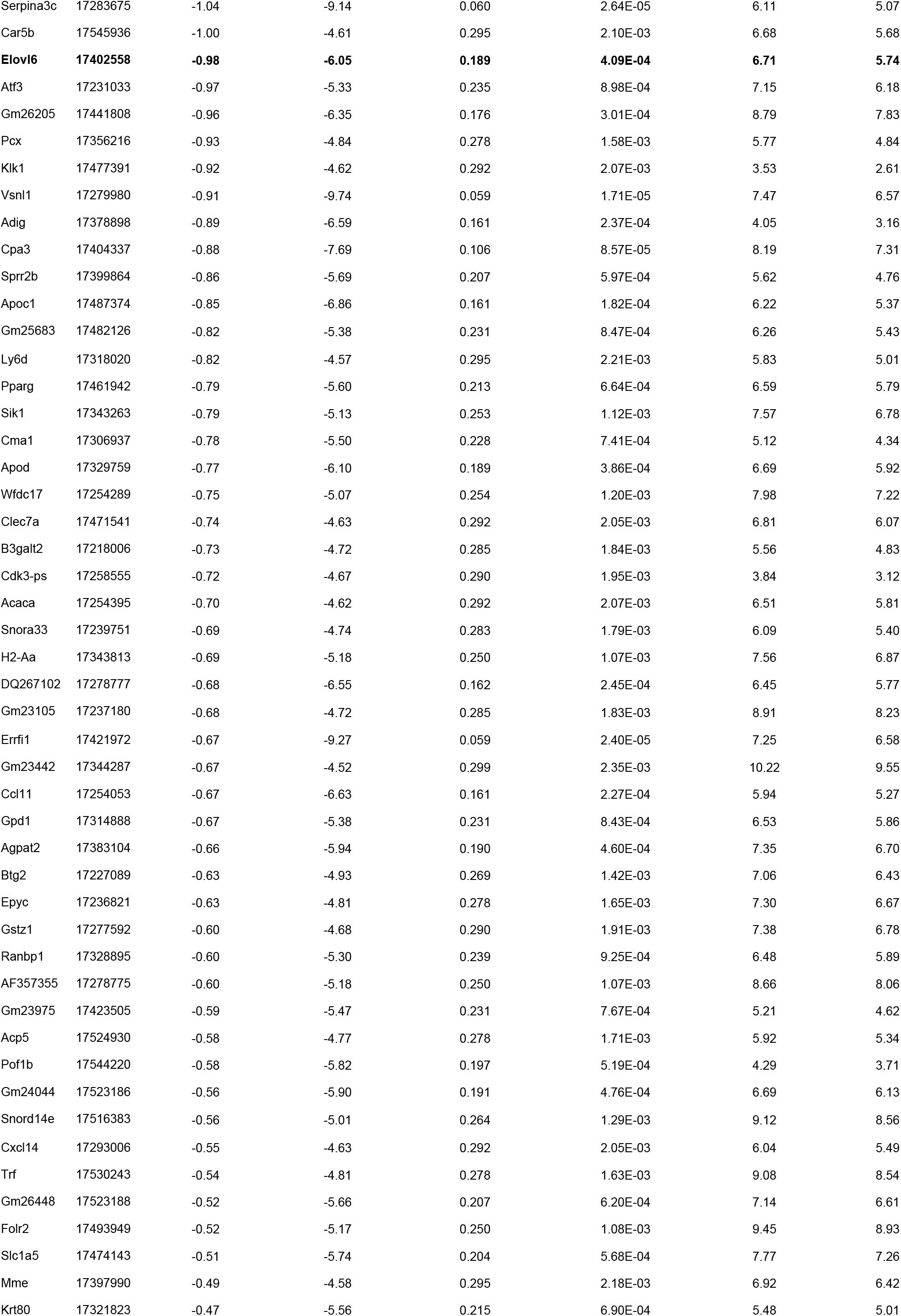
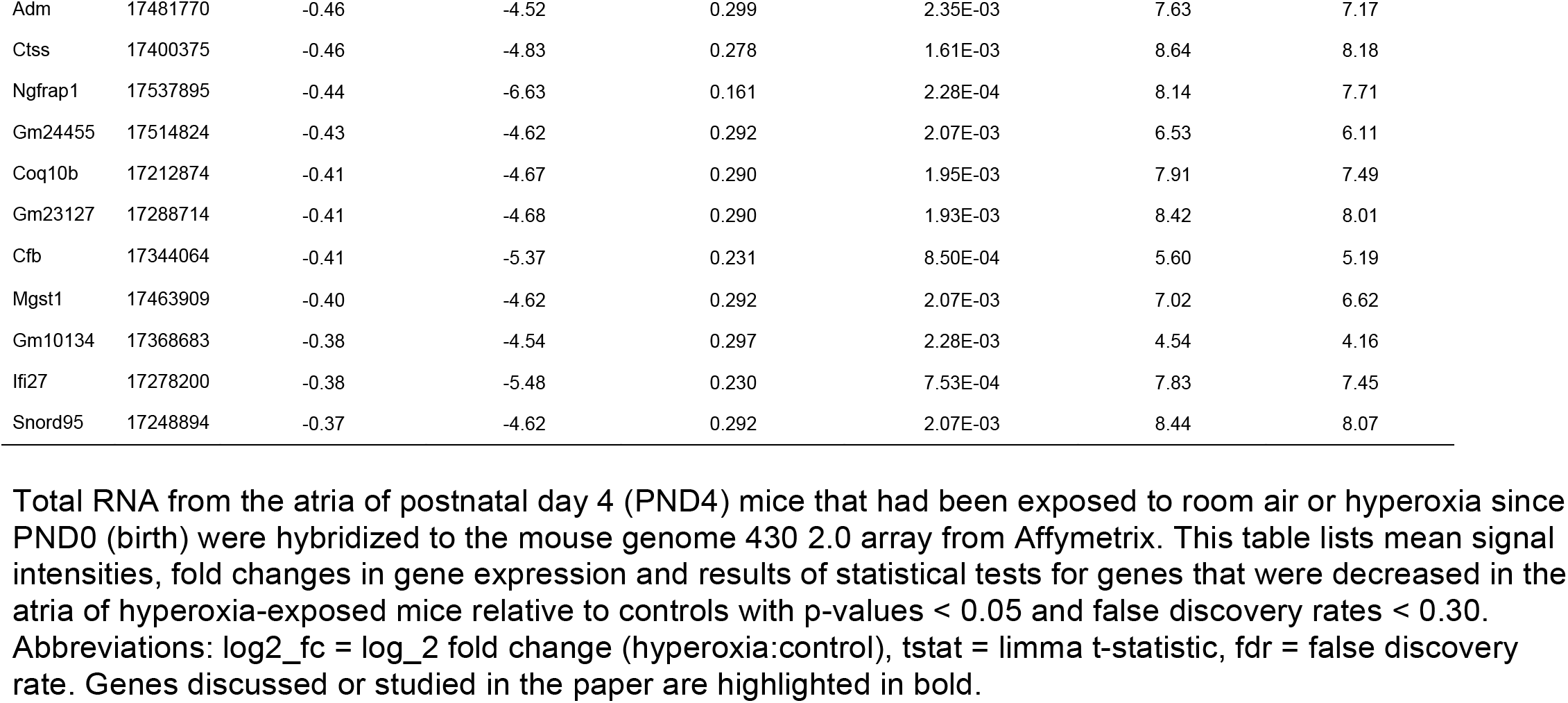
Genes expressed at lower levels in the atria of neonatal hyperoxia-exposed mice relative to controls.

**Supplemental Table 4.**
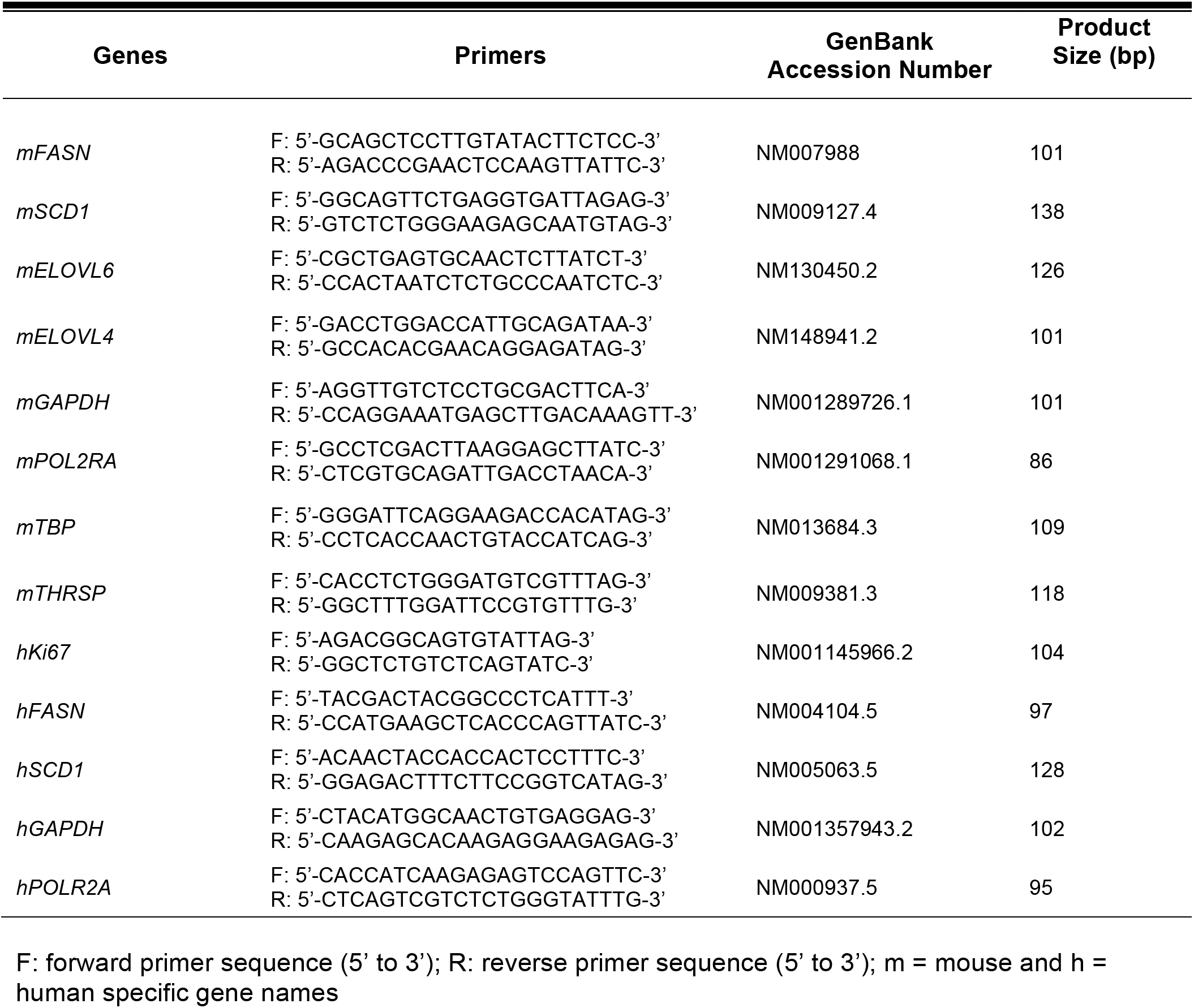
List of primers used for qRT-PCR.

